# Heterogeneity of normal human breast stem and progenitor cells as revealed by transcriptional profiling

**DOI:** 10.1101/109751

**Authors:** Justin A. Colacino, Ebrahim Azizi, Michael D. Brooks, Shamileh Fouladdel, Sean P. McDermott, Michael Lee, David Hill, Maureen A. Sartor, Laura S. Rozek, Max S. Wicha

## Abstract

During development and pregnancy, the human mammary gland undergoes extensive remodeling in processes driven by populations of stem and progenitor cells. We recently reported that breast cancers are also hierarchically organized and driven by distinct populations of cancer stem cells characterized as CD44^+^CD24^low/−^ or by expression of Aldehyde dehydrogenase (ALDH). These sets of markers identify largely non-overlapping mesenchymal and epithelial populations, each of which is capable of tumor initiation when transplanted into immunosuppressed mice. Less is known about these two populations, individually or their overlap, in the normal human mammary gland. The goal of this study was to understand the biology of the ALDH^+^ and CD44^+^CD24^−^ populations in the normal human breast, using flow cytometry based sorting paired with functional *ex vivo* analyses, RNA-sequencing, and single cell RNA expression profiling. ALDH^+^ cells and ALDH^−^CD44^+^CD24^−^ cells, generally, have epithelial-like and mesenchymal-like characteristics, respectively. Despite this, there are substantial similarities in the biological pathways activated in both populations when compared to differentiated cells. Additionally, we found a substantial proportion of cells that simultaneously express ALDH^+^ and CD44^+^CD24^−^ whose abundance varies between individuals. At the single cell level, these cells have the greatest mammosphere forming capacity and express high levels of stemness and EMT-associated genes including *ID1, SOX2, TWIST1, and ZEB2.* Through unbiased analysis of individual ALDH+ cells, we find cells with either epithelial or mesenchymal expression phenotypes. We also identify a subpopulation of cells with a hybrid epithelial/mesenchymal expression phenotype that overexpress genes associated with aggressive triple negative breast cancers. These results highlight the utility of single cell analyses to characterize tissue heterogeneity, even in marker enriched cell populations, and further identifies the genes and pathways that define this heterogeneity.

## Introduction

*In utero,* throughout puberty, and during pregnancy, the human mammary gland undergoes extensive expansion and remodeling, driven by populations of stem and progenitor cells (Visvader and Stingl, 2014). The mammary epithelium consists of two major cellular lineages, luminal and myoepithelial. Lineage tracing studies have shown that the mammary differentiation hierarchy consists of bipotent mammary stem cells which can give rise to both luminal and myoepithelial cells (Rios et al., 2014), as well as long lived unipotent progenitor cells which drive mammary gland development and homeostasis (Van Keymeulen et al., 2011). Breast cancers also display a differentiation hierarchy and are driven and sustained by a stem -like cell population (Al-Hajj et al., 2003). The long-lived nature and the proliferative capacity of bipotent breast stem cells or lineage committed progenitor cells make these good candidates to be the cells of origin of breast cancers. Molecular analysis of breast cancers has led to identification of subtypes with distinct gene expression profiles and clinical behaviors. This has led to the hypothesis that different breast tumor subtypes arise from cells in the normal breast hierarchy (Skibinski and Kuperwasser, 2015). Despite substantial advances in the field, the cells of origin of different subtypes of human breast cancers have not been conclusively identified and it remains unclear whether the differentiation hierarchy in breast cancers reflects that in the normal human breast.

We have reported that aldehyde dehydrogenase activity (termed ALDH^+^) and CD44^+^CD24^−^ mark two largely non-overlapping populations of cancer stem cells, which have epithelial- and mesenchymal-like phenotypes, respectively (Liu et al., 2013). These cells are endowed with plasticity allowing them to inter-convert, in a process driven by the tumor microenvironment. This plasticity may play an important role in the successful execution of the metastatic process (Liu et al., 2013). The existence of these two stem cell populations mirrors that of the long-lived unipotent progenitor populations in the normal breast. The markers ALDH^+^ (Ginestier et al., 2007) and CD44^+^CD24^−^ (Choudhury et al., 2013; Shipitsin et al., 2007) have also proven useful in the isolation and functional characterization of normal human breast stem cells. A number of other markers, such as CD49f^hi^EpCAM^−/lo^ (Stingl et al., 2001; Visvader, 2009) have also been reported to enrich for normal human breast stem cells. The importance of the ALDH^+^ and CD44^+^CD24^−^ cell populations in breast cancer, and the comparatively little knowledge about cells that express either or both markers in the normal breast highlight the importance of further charactering these populations.

The goal of this study was to understand the biology of the ALDH^+^ and CD44^+^CD24^−^ populations in the normal human breast, using flow cytometry based sorting paired with functional *ex vivo* analyses, RNA-sequencing, and single cell RNA profiling. Unlike in breast cancers, we identified a significant overlap between the ALDH^+^ and CD44^+^CD24^−^ populations in the normal breast, with substantial interdindividual variation in the degree of overlap. While ALDH^+^ cells and ALDH^−^CD44^+^CD24^−^ (hereto referred to as CD44^+^) cells, generally, represent epithelial-like and mesenchymal-like breast populations, there are substantial similarities in the biological pathways activated in both populations when compared to differentiated ALDH^−^CD44^−^CD24^+^ (hereto referred to as CD24^+^) cells. The cells that express both ALDH^+^ and CD44^+^CD24^−^ have the greatest mammosphere formation potential, and express higher levels of stemness and epithelial-to-mesenchymal transition (EMT)-related genes at both the population and single cell level. By conducting an unbiased analysis of single cells, we identified a substantial degree of cellular heterogeneity within the ALDH+ and CD44+/CD24-cell populations. In addition, we demonstrate the existence of a subpopulation of ALDH^+^ cells within the normal breast that simultaneously express both epithelial and mesenchymal markers. Expression of these markers is associated with poor outcome in a cohort of patients with triple negative breast cancer.

## Materials and Methods

### Tissue Procurement

Normal (non-pathogenic) breast tissue was isolated from women undergoing voluntary reduction mammoplasty at the University of Michigan. The study protocol was approved by the University of Michigan Institutional Review Board. Breast tissue was mechanically and enzymatically digested as previously described (Dontu et al., 2003; Kakarala et al., 2010).

### Flow Cytometry

Normal mammary cells were stained for CD44, and CD24 expression and ALDH activity accessed by the Aldefluor assay as described previously (Colacino et al., 2016) and analyzed or sorted on a Beckman Coulter MoFlo Astrios at the University of Michigan Flow Cytomtery core facility. Briefly, single mammary cells were first incubated with a lineage depletion cocktail that consisted of biotinylated antibodies targeted against CD45, HLA-DR, CD14, CD31, CD41, CD19, CD235a, CD56, CD3, CD16, and CD140b (all from eBioscience, except for CD140b (Biolegend) and CD41 (Acris)). Next, cells were stained with Alexafluor 750 tagged streptavidin, LIVE/DEAD Fixable Dead Cell Stain (Invitrogen), CD24 (Biolegend), CD44 (BD), and Aldefluor (Stem Cell Technology). Single color and isotype controls were included for compensation and gating purposes. Aldefluor-positive gating was based on DEAB (negative) controls. Flow cytometry data analysis was performed with FlowJo software v10.0.8.

### RNA Extraction and Sequencing

ALDH^+^, CD44^+^, and CD24^+^ cell populations were collected as described above and total RNA was isolated from each cell population using the RNEasy Micro Kit (Qiagen) with on column DNase treatment. RNA concentration and quality was determined using a Nanodrop (Thermo) and Bioanalyzer (Agilent). Ribosmal RNAs were depleted using Ribominus (Life) and sequencing libraries were prepared with the SMARTer Stranded RNA-Seq kit (Clontech). Libraries were multiplexed (4 per lane) and sequenced using paired end 50 cycle reads on a HiSeq 2500 (Illumina) at the University of Michigan DNA Sequencing Core Facility.

### RNA-Seq Data Analysis

The Flux high-performance computer cluster hosted by Advanced Research Computing (ARC) at the University of Michigan was used for computational analysis. Sequencing read quality was assessed utilizing FastQC. Sequencing reads were concatenated by sample and read in pair using SeqTK. The first three nucleotides of the first read in each read pair were trimmed, as recommended by Clontech, using Prinseq 0.20.3. A splice junction aware build of the human genome (GRCh37) was built using the genomeGenerate function from STAR 2.3.0 (Dobin et al., 2013). Read pairs were aligned to the genome using STAR, using the options “outFilterMultimapNmax 10” and “sjdbScore 2”. The aligned reads were assigned to genomic features (GRCh37 genes) using HTSeq-count, with the set mode “union”. We conducted differential expression testing on the assigned read counts per gene utilizing edgeR (Robinson et al., 2010). Separate comparisons were conducted for each cell type (ALDH^+^, CD44^+^, and CD24^+^), adjusting for study subject as a covariate using glmLRT. To reduce the dispersion of the dataset due to lowly expressed genes, genes with a mean aligned read count less than five across all samples were excluded from analysis. Normalized counts per million were estimated utilizing the “cpm” function in edgeR (Robinson et al., 2010). Genes were considered differentially expressed between cell populations at a false discovery rate (FDR) adjusted p-value < 0.05 (Benjamini and Hochberg, 1995).

### Pathway Analyses, Integration with Publically Available Data

Differentially expressed pathways were identified utilizing iPathwayGuide (AdvaitaBio). A directional analysis was conducted on all genes by including p-value of the differential expression test between groups as an effect size measure and log2 fold difference in expression as a measure of effect direction. Biological pathways were considered differentially expressed at a FDR P-value <0.05. To compare how genes identified as differentially expressed between the ALDH^+^ and CD44^+^ cells overlap with previously reported breast stem and progenitor cell gene expression signatures, we compared log fold changes in expression in our data for genes identified as uniquely upregulated (logFC >1) in CD49f+/EpCAM- (“mammary stem”) and CD49f+/EpCAM+ (“luminal progenitor”) cell populations (Lim et al., 2009). Comparisons of the Cancer Genome Atlas expression data between triple negative breast cancers and other cancers were conducted using Oncomine (https://www.oncomine.org/resource/login.html). Estimation of recurrence free survival differences by expression of specific genes was conducted using KMPlot (Gyorffy and Schafer, 2009) for four gene probes: 209086_x_at (*CD146*), 203237_s_at (*NOTCH3*), 213342_at (*YAP1*), 214031_s_at (*KRT7*). Survival analyses were restricted to ER negative, PR negative, HER2 tumors (n=249 total patients), with groups dichotomized using the auto selected best cutoff. Additionally, survival based on the mean expression of these four probes was also estimated.

### Mammosphere formation

Single cells were sorted from reduction mammoplasty tissue from three individuals by flow cytometry as described above and plated in 96 well ultralow attachment plates (Corning) at a density of 500 cells per well. Four breast cell populations were sorted and plated: ALDH^+^CD44^+^CD24^−^, ALDH^−^CD44^+^CD24^−^, ALDH^+^ that are not CD44^+^CD24^−^, and ALDH- cells that are not CD44^+^CD24^−^. Mammospheres formed for 7-10 days in Mammocult media (StemCell). Each experiment was run with at least three replicates and primary sphere number was quantified manually.

### Single Cell Transcriptional Profiling

Single ALDH^+^CD44^+^CD24^−^ and ALDH^+^ that are not CD44^+^CD24 mammary epithelial cells sorted by flow cytometry (explained above), were stained with CellTracker dyes to differentiate the two populations and loaded onto the Fluidigm’s C1 PreAmp chip to isolate up to 96 single cells. C1 chips were examined under the Olympus IX82 invert fluorescent microscope to verify chambers contain single cells. The captured cells in each chamber of the C1 chip then sequentially underwent lysis, RNA release, cDNA synthesis and preamplification of 96 target genes. The preamplified cDNAs were then analyzed using Fluidigm’s Biomark HD system, 96x96 chip, and 96 TaqMan assays to determine expression patterns of 96 target genes in each cell. Finally, the qPCR data were analyzed using Fluidigm’s R Package SINGuLAR to generate violin, heatmap clustering and principal component analysis (PCA) plots based on the Log2 expression values for each gene in each cell.

## Results

### Isolation and Characterization of Normal Mammary Cell Populations

To follow up on our findings of epithelial-like and mesenchymal-like breast cancer stem cells (Liu et al., 2013), we isolated three cellular populations from the normal breast by flow cytometry: ALDH^+^, CD44^+^, and CD24^+^ (Figure 1A). Through RNA-sequencing, we confirmed that the expression of *ALDH1A1*, *ALDH1A3*, *CD44*, and *CD24* matched the protein markers used for sorting (Figure 1B). Multidimensional scaling identified that the samples cluster on the first two dimensions of the leading log fold change, with the ALDH^+^ and the CD24^+^ cells grouping together on the first dimension, but separating on the second (Figure 1C). Differential expression analysis identified broad gene expression differences between the populations (Figure1D).

**Figure 1.**
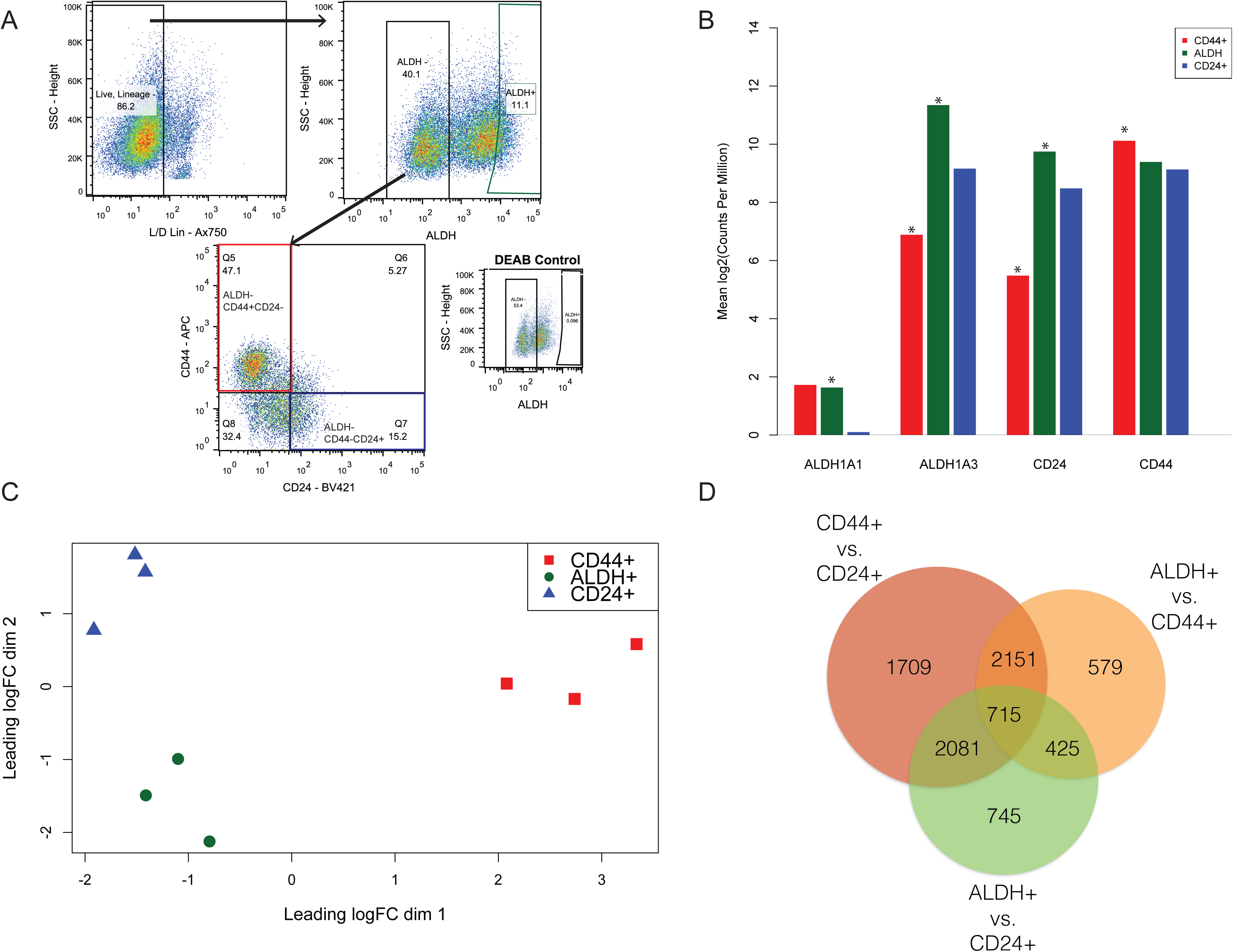
Purification and transcriptomic profiling of ALDH^+^, ALDH^−^CD44^+^CD24^−^, and ALDH^−^CD44^−^CD24^+^ normal human breast cells. (A) A representative FACS isolation diagram of the three cellular populations from normal mammary cells isolated from voluntary reduction mammoplasty tissues. ALDH+ gating was based on the DEAB negative control. ALDH^−^CD44+CD24-will be hereto referred to as CD44^+^ and ALDH^−^CD44^−^CD24^+^ as CD24^+^ (B) RNA expression, from RNA-seq analysis, of genes associated with the markers used for population sorting. * -FDR p <0.05 (C) Multidimensional scaling plot of the three cellular populations based on the 500 most variably expressed genes. (D) Overlap in differentially expressed (FDR p < 0.05) genes between the three populations.

### The ALDH^+^ Breast Cell Gene Expression Signature

We have previously shown that normal breast and breast cancer cells with high aldehyde dehydrogenase activity (ALDH^+^) are enriched for stem-like cells (Ginestier et al., 2007). To quantify expression patterns specific to normal breast ALDH^+^ cells, we compared expression of ALDH^+^ cells to CD24^+^ cells, which do not express either of the canonical breast stem cell markers ALDH or CD44^+^CD24^−^. In ALDH^+^ cells, 2244 genes were upregulated and 1730 downregulated (Figure 2A; **Supplemental Table 1**). The top 3 most overexpressed genes, by magnitude of effect were *WNT2* (fold change = 705.3), Insulin-like growth factor 1 (*IGF1;* fold change = 532.3), and the canonical notch ligand *DLL1* (fold change = 502.5). Analyzing the relative expression of other WNT pathway genes showed that, in addition to *WNT2*, ALDH^+^ cells also overexpress *RSPO3*, *SFRP4*, *MMP7*, and *FOSL1* (Figure 2D). We next compared the ALDH+ cell expression signature to previously reported gene expression signatures of human mammary stem (CD49f^+^/EpCAM-) and luminal progenitor (CD49f^+^/EpCAM^+^) cells (Lim et al., 2009). We did not observe a strong enrichment for either the mammary stem or luminal progenitor gene signature in ALDH^+^ cells (Figures 2B-2C). KEGG pathway analyses identified that ALDH^+^ cells differentially expressed genes involved in ribosome (FDR=3.1E-16), oxidative phosphorylation (FDR=2.6E-14), and the proteasome (7.2E-14) (**Supplemental Figures 1-3**). In each of these three pathways, all of the differentially expressed genes were upregulated in the ALDH^+^ cells. Genes differentially expressed in ALDH^+^ cells were also enriched in pathways related to focal adhesion (p=1.9E-4) and ECM-receptor interactions (p=9.8E-7) (**Supplemental Table 2**).

**Figure 2.**
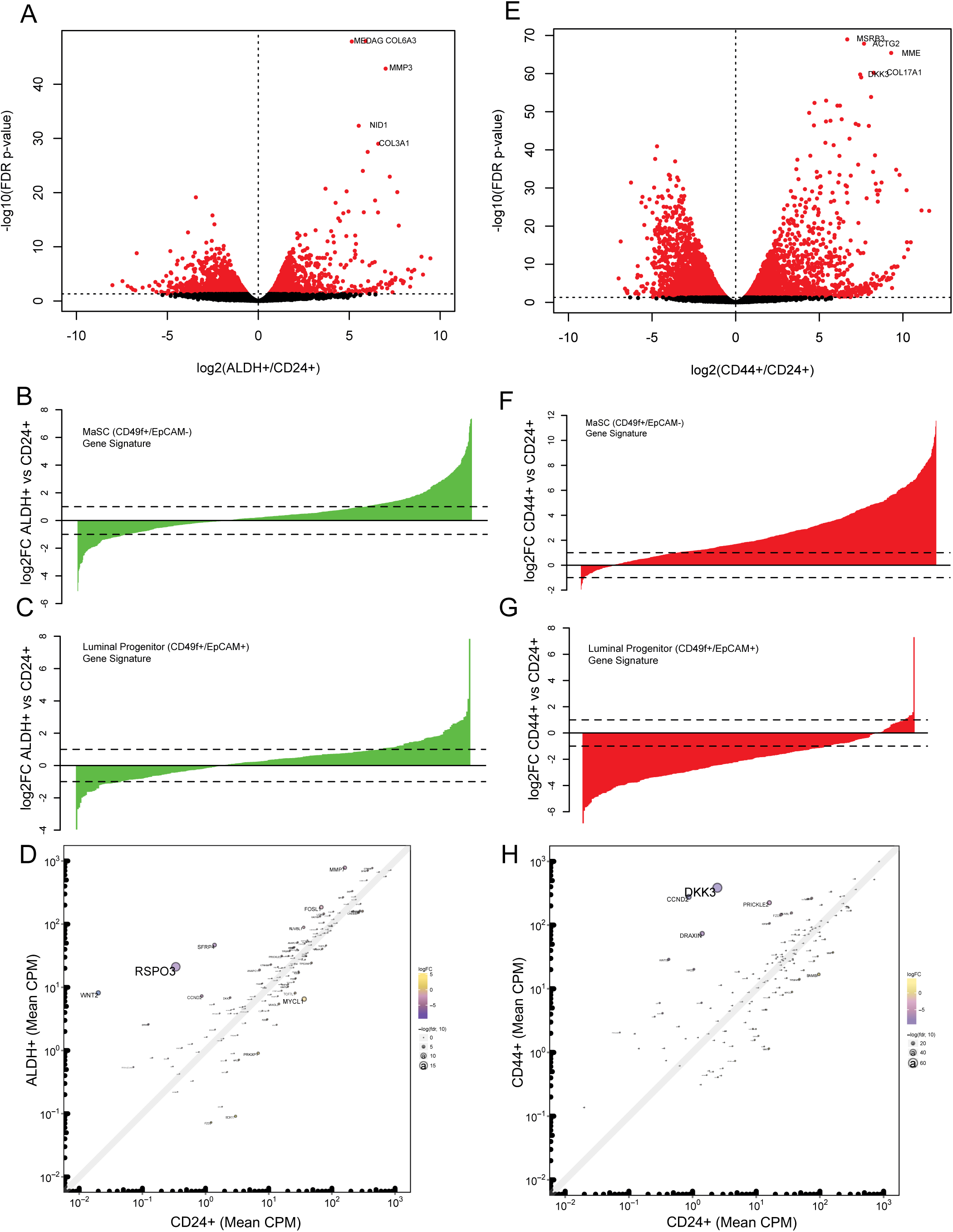
Comparison of gene expression signatures between ALDH^+^ and ALDH^−^CD44^+^CD24^−^ with non-stem cell enriched ALDH^−^CD44^−^CD24^+^ cells. (A) FDR volcano plot comparing the log2-fold change in gene expression between ALDH+ and CD24^+^ cells, with the names of the top 5 most statistically different genes labeled. (B and C) Comparison of the log2-fold change differences between ALDH+ and CD24^+^ and the mammary stem cell and luminal progenitor cell gene expression signature, respectively, reported in Lim et al., 2009. (D) Enrichment of Wnt signaling related genes in ALDH+ relative to CD24^+^ cells. (E) FDR volcano plot comparing differences in gene expression between CD44+ and CD24^+^ cells. (E and F) Comparison of the log2-fold change differences between CD44+ and CD24^+^ and the mammary stem cell and luminal progenitor cell gene expression signature, respectively. (H) Enrichment of Wnt signaling related genes in CD44+ relative to CD24^+^ cells.

### The ALDH^−^CD44^+^CD24^−^ Gene Expression Signature

We further compared the gene expression profile of CD44^+^ cells to CD24^+^ cells, and identified 3361 genes statistically significantly upregulated and 3295 genes downregulated (Figure 2E, **Supplemental Table 3**). The most overexpressed genes by magnitude of effect in the CD44^+^ cells were Insulin-like Growth Factor Binding Protein 1 (*IGFBP1*; fold change = 3040.3), the glycosyltransferase *ST8SIA2* (fold change = 2,210.3), phospholipase D family member 5 (*PLD5*, fold change = 1418.4), the chaperone protein *SCG5* (fold change = 1269.5), and the cytoskeletal protein *MYOT* (fold change = 1209.3).

These genes had no detectable expression in CD24^+^ cells (**Supplemental Table 3**). The expression pattern of Wnt pathway members in CD44^+^ cells was substantially different than for ALDH^+^ cells, with the signature of CD44^+^ cells driven by *DKK3*, *CCND2*, *PRICKLE2*, and *DRAXIN* (Figure 2H). CD44^+^ cells had enrichment for the previously reported mammary stem (CD49f^+^/EpCAM-) gene signature (Figure 2F), and negative enrichment for the luminal progenitor signature (Figure 2G). KEGG pathway analysis found that CD44^+^ cells overexpress genes associated with the proteasome (FDR=3.8E-8) and ECM-Receptor interactions (FDR=5E-6), as well differentially express a suite of genes associated with Focal Adhesion (FDR=1.6E-6) (**Supplemental Figures 4-6**). Further, CD44^+^ cells differentially expressed genes involved in PI3K-AKT signaling (FDR=7.0E-5), including *CCDN2*, *PIK3CG*, *FGF1*, and *NGF* (**Supplemental Figure 7**).

### CD44^+^ Cells Express Mesenchymal Markers, whereas ALDH^+^ Cells Express Both Epithelial and Mesenchymal Markers

We have previously shown that ALDH^+^ cells cultured as mammospheres for 24 hours had a global gene expression pattern reflecting an epithelial phenotype relative to CD44^+^ cells, which have a generally mesenchymal phenotype (Colacino et al., 2016). To further explore the relative phenotypes of the ALDH^+^, CD44^+^, and CD24^+^ cells, we compared the expression of genes associated with an epithelial or mesenchymal phenotype across these populations (Figure 3). In general, we observed that CD44^+^ were strongly enriched for expression of mesenchymal markers, including *CDH2*, *KRT17*, *ZEB2*, *KLF8*, *CD44*, *KRT14*, and *VIM*. ALDH^+^ cells expressed the highest levels of various epithelial markers, including *KRT19*, *CD24*, *CDH1*, *EpCAM*, and *KR18*. ALDH^+^ cells also, however, expressed intermediate or highest levels of several mesenchymal markers, including the EMT transcription factors *SNAI1, TWIST1*, and *ZEB2*. These results suggest that ALDH^+^ cells are either simultaneously expressing both epithelial and mesenchymal markers, or that there are epithelial-like and mesenchymal-like subpopulations within the ALDH^+^ cell fraction.

**Figure 3.**
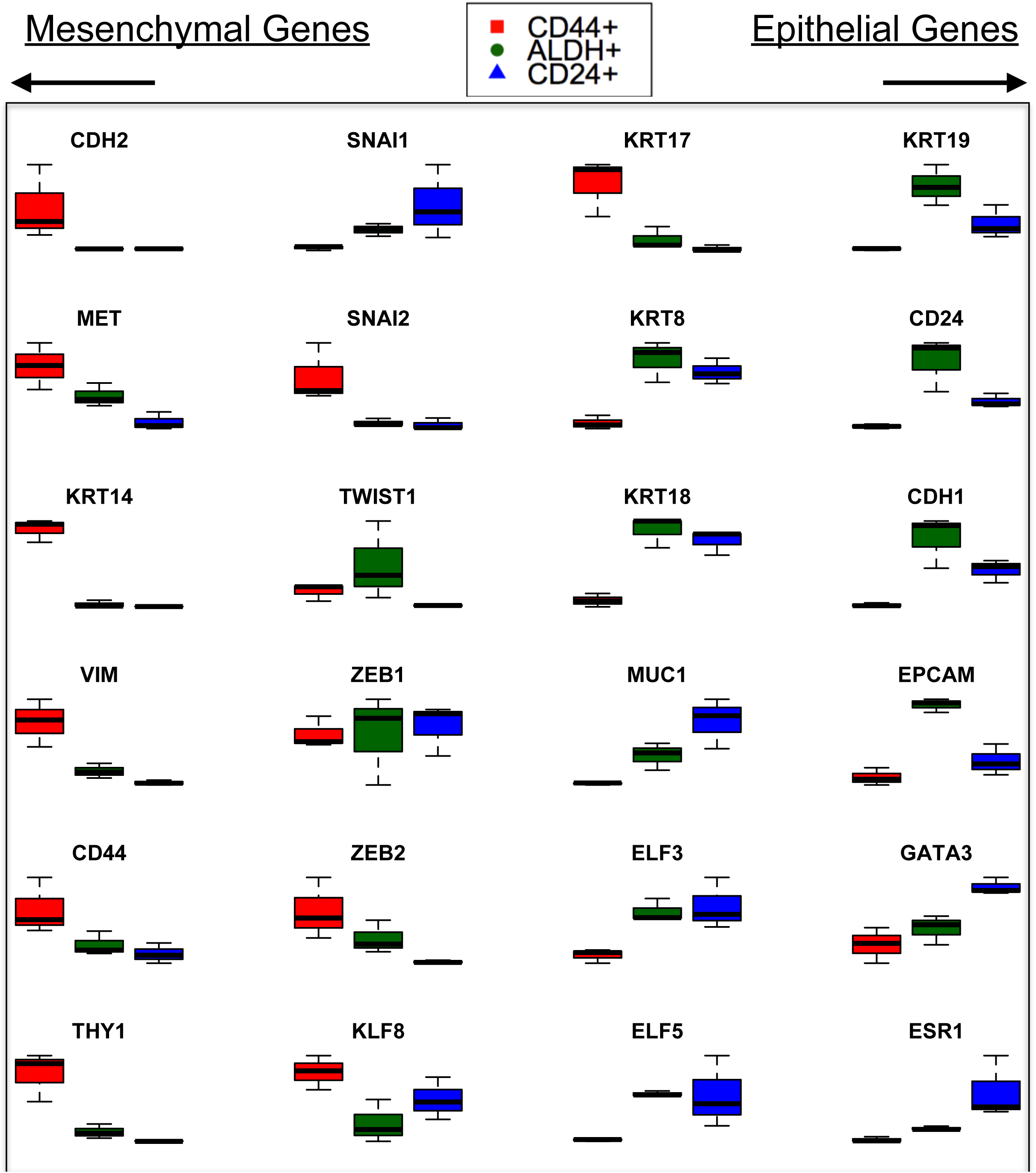
Relative expression levels of mesenchymal phenotype-and epithelial phenotype-associated genes in ALDH^+^, CD44^+^, and CD24^+^ cells.

### ALDH^+^ Cell Populations Are Highly Variable Between Individuals

To better understand the heterogeneity within the ALDH^+^ normal mammary cell population, we conducted flow cytometry analysis of ALDH activity combined with CD44 and CD24 staining across samples obtained from 8 women. We identified significant inter-individual variation in the proportion of ALDH^+^ cells (range 8.9% - 45%) (Figure 4A). When analyzing the relative proportion of CD44^+^CD24^−^ expressing cells within the ALDH^+^ fraction, we identified further heterogeneity between individuals. The proportion of CD44^+^CD24^−^ cells within the ALDH^+^ fraction ranged from 13.3% to 70.3%. The majority of ALDH^+^ cells tended to also be CD44^+^. To understand the functional differences between these different cell populations in the normal human breast, we isolated 4 distinct populations: ALDH^+^CD44^+^CD24^−^ cells, ALDH^+^ cells that are not CD44^+^CD24^−^, ALDH^−^CD44^+^CD24^−^ cells, and ALDH^−^ cells that are not CD44^+^CD24^−^ and determined their capacity to form mammospheres a property associated with “stemness”. ALDH^+^CD44^+^CD24^−^ cells displayed the highest capacity to form mammospheres. Both ALDH^+^ cells that are not CD44^+^CD24^−^ and ALDH^−^ CD44^+^CD24^−^ also formed mammospheres, but at rates less than ALDH^+^CD44^+^CD24^−^ cells (Figure 4B). ALDH^−^ cells that are not CD44^+^CD24^−^ cells did not form mammospheres. These results suggest that the majority of the cells with mammosphere forming potential in the normal human breast lie in the ALDH^+^CD44^+^CD24^−^ cell fraction.

**Figure 4.**
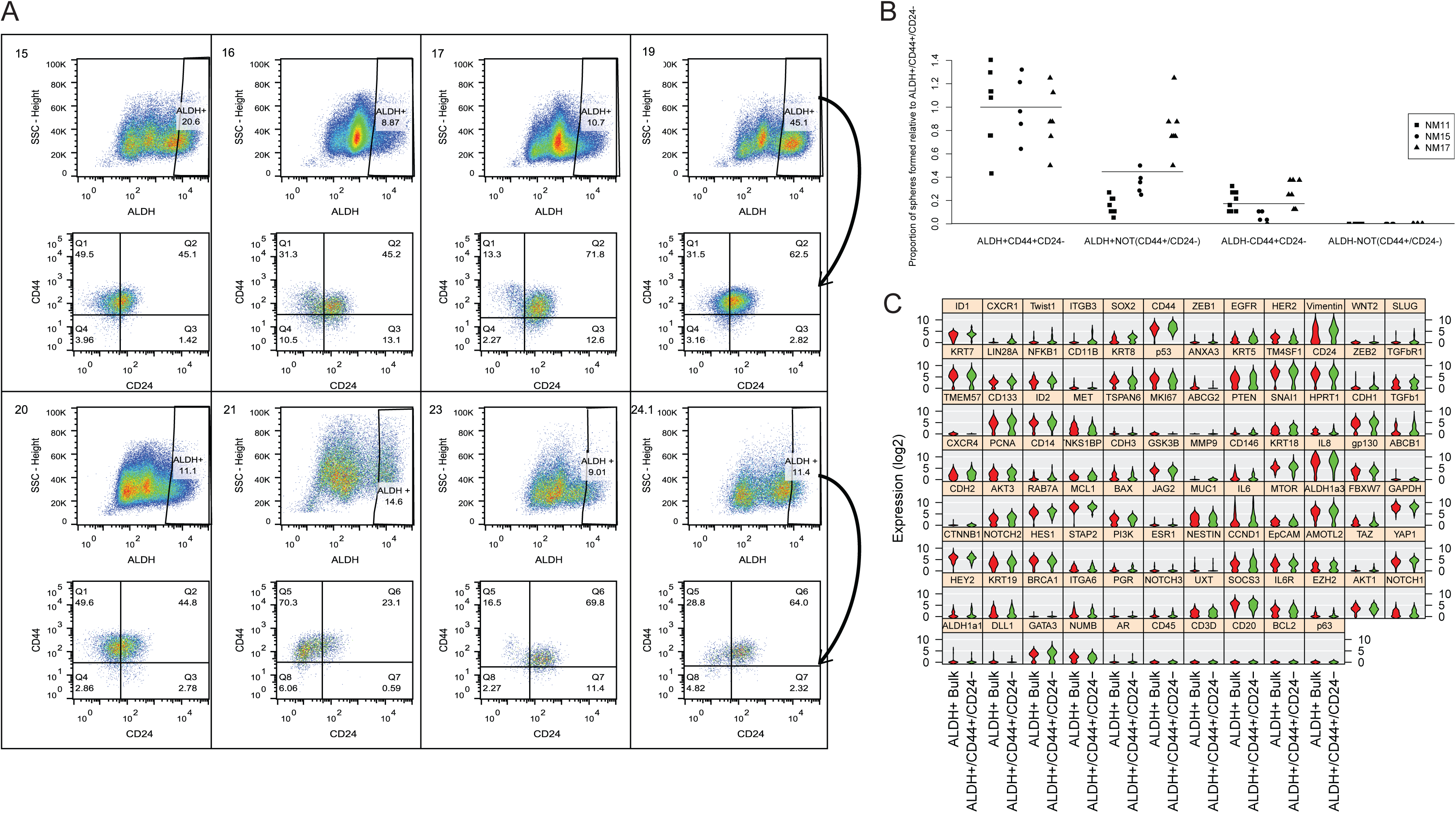
Quantitation and profiling of normal breast cells that express both stem cell markers ALDH^+^and CD44^+^CD24^-^. (A) Quantitation, by flow cytometry, of the ALDH^+^cell population in normal mammary tissues (n=8). ALDH^+^ cells were then further analyzed for CD44 and CD24 expression (arrow), with the top left quadrant in each bottom panel representing ALDH^+^CD44^+^CD24^−^ cells. (B) Mammosphere formation rates of cell populations expressing different combinations of ALDH and CD44/CD24 (cells sorted from n=3 individuals). (C) Single cell gene expression profiling of ALDH^+^CD44^+^CD24^−^ and ALDH^+^ Bulk (ALDH^+^ cells which do not express CD44^+^CD24^−^) cells. Genes are ordered by statistical significance of difference in expression between the two populations.

### Single Cell RNA Profiling Reveals Further Heterogeneity Within the ALDH^+^ Cell Population

To characterize the expression differences, and the expression heterogeneity, that define ALDH^+^CD44^+^CD24^−^ cells and the remainder of ALDH^+^ cells we isolated these two fractions by FACS, fluorescently labeled each population, and subjected them to both bulk RNA and single cell RNA expression analysis of a custom 96 gene panel using Fluidigm’s C1 and Biomark HD instruments. This panel contains genes previously demonstrated to be expressed in stem cells or in regulating EMT (Holm et al., 2010; Jolly et al., 2015; Liu et al., 2013; The Cancer Genome Atlas, 2012). Utilizing the RNA-seq data described above, we confirmed that the custom panel effectively discriminated ALDH^+^, CD44^+^, and ^−^CD24^+^ cells populations by MDS (**Supplemental Figure 8A**).

Comparing RNA isolated from bulk ALDH^+^CD44^+^CD24^−^ and ALDH^+^ cells that are not also CD44^+^CD24^−^, we identified that ALDH^+^CD44^+^CD24^−^ cells expressed lower levels of *BAX*, *CDH3*, *CDH1*, *KRT19*, and *MUC1* (p<0.10; **Supplemental Figure 8B**; **Supplemental Table 4**). To characterize the heterogeneity within these sorted populations we analyzed expression of the 96-gene panel in single cells isolated by Fluidigm C1 technology. When comparing gene expression profiles at the single cell level to those of bulk cells, a different pattern emerged (Figure 4C). At the single cell level, ALDH^+^CD44^+^CD24^−^ cells overexpress the stem cell genes *ID1* and *SOX2*, the IL-8 receptor *CXCR1*, and the EMT-associated transcription factors *TWIST1* and *ZEB2.* ALDH^+^CD44^+^CD24^−^ cells also significantly overexpressed *EGFR, CD44, WNT2*, and *VIM*, while expressing *HER2* at lower levels. These results illustrate the heterogeneity within the ALDH+ population and highlight that there are subpopulations of ALDH^+^ cells with a range of cellular phenotypes. Additionally, these results show the importance of analyzing single cells, even within flow cytometry sorted populations.

To further explore the heterogeneity of ALDH^+^ cells, we conducted unsupervised hierarchical clustering of the gene expression patterns of single ALDH^+^ cells (n=105 total from 3 individuals). We observed four clusters, presented by heatmap (Figure 5A) and PCA analysis (Figure 5B). Comparing the gene expression patterns for the four clusters across the 96 genes on our panel revealed substantial differences, with 75 of the genes being significantly different across the groups by ANOVA (p<0.05; Figure 5C; **Supplemental Table 5**). Cluster 1 was characterized, in general, by low overall gene expression. Cluster 4 expressed high levels of epithelial-related genes, including *KRT7*, *CD24, EPCAM*, *GATA3*, and *KRT5*. Cluster 3 was characterized by expression of mesenchymal-phenotype related genes, including *CD44*, *ZEB2*, *SLUG*, *VIM*, *TWIST1*, *ZEB1*, and *TGFB1*. Cluster 2, which had the highest average expression of *ITGA6, CD133,* and *ALDH1A3*, expressed high levels of epithelial-related genes, such as *KRT7, KRT8*, and *CDH1* as well as genes associated with EMT, including *IL6*, *CD44*, *TM4SF1*, and *VIM*, and the mesenchymal stem cell marker *CD146*. Cells in this cluster express canonical markers of breast stemness at the highest levels and present with a hybrid epithelial/mesenchymal phenotype (Figure 5D). Cluster membership was found to be strongly associated with the patient from which the cells were isolated (Figures 5E and 5F), with themajority of Cluster 2 cells belonging to patient NM15. This emphasized the variability of these cell population in different individuals.

**Figure 5.**
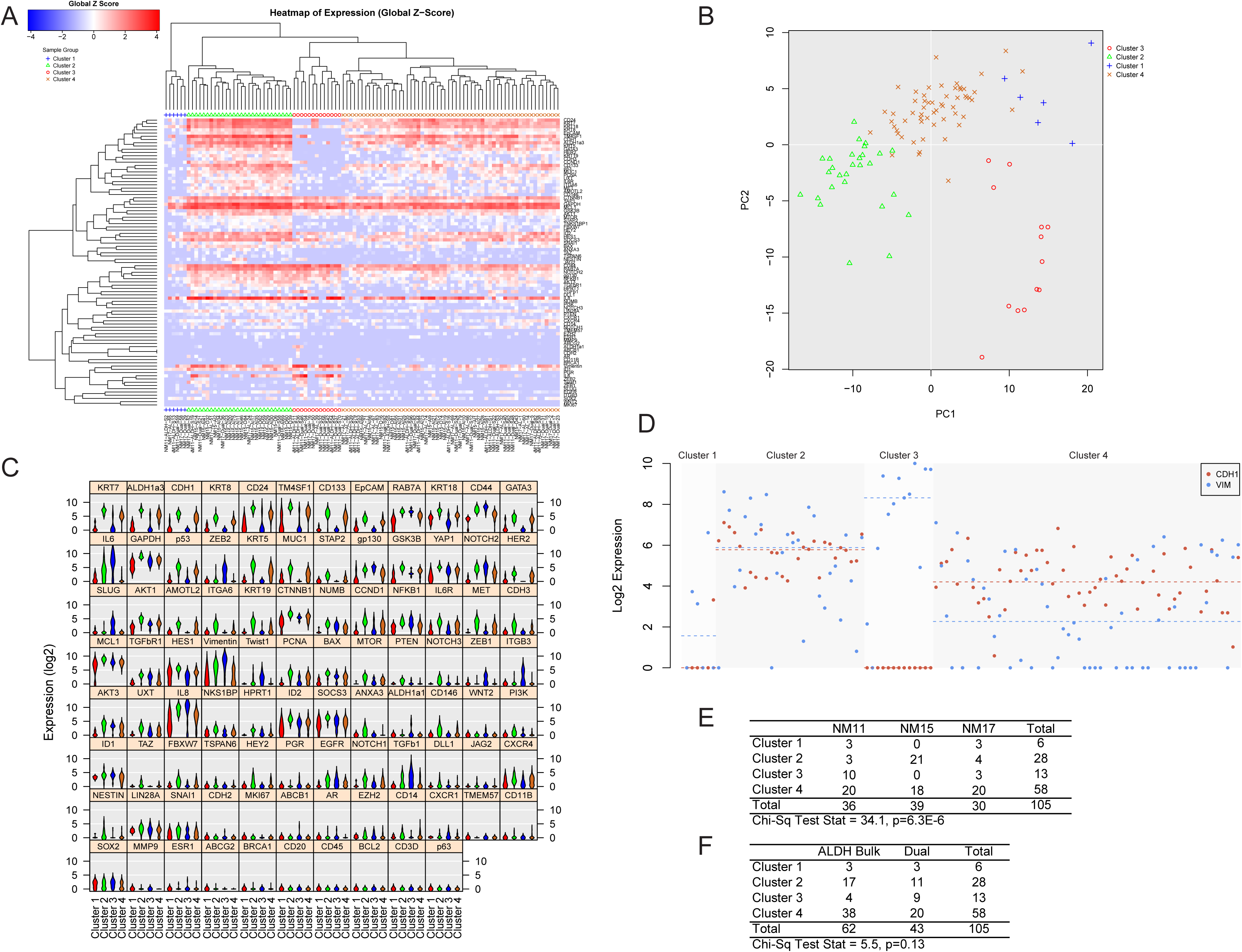
Unbiased analysis of single cell expression data from ALDH+ normal breast cells. (A) Hierarchical clustering analysis of gene expression measures from 105 ALDH+ cells from 3 individuals reveals 4 expression clusters. (B) Principal component analysis of the gene expression profiles of the 105 ALDH+ cells. Violin plot analysis of the expression of the targeted gene expression panel across the four clusters. (D) Comparison of expression of the epithelial marker gene *CDH1* (red) and the mesenchymal marker gene *VIM* (blue) across cells in the four clusters. (E) Distribution of the four expression clusters across the cells isolated from the three study subjects. (F) Distribution of ALDH^+^CD44^+^CD24^−^ and ALDH^+^ Bulk cells across the four expression clusters.

### ALDH1A1 Expressing Cells Have a Mesenchymal-Like Phenotype

Cells in Cluster 3 expressed the highest levels of *ALDH1A1* and low levels of *ALDH1A3*, while cells in Clusters 2 and 4 expressed high levels of *ALDH1A3*, but low levels of *ALDH1A1*. This segregated expression of *ALDH1A1* and *ALDH1A3* corroborates previous reports of differential localization of ALDH1A1 and ALDH1A3 protein expression in the normal mammary gland (Honeth et al., 2015). To further explore these differences, we compared the gene expression patterns between *ALDH1A1* expressing cells and the remainder of the ALDH^+^ cells (**Supplemental Figure 9**). *ALDH1A1*^*+*^ cells expressed significantly higher levels of mesenchymal associated markers, including *TWIST1*, *ZEB2*, *IL6*, *ZEB1*, *VIM*, and *TGFB1* and lower levels of epithelial associated markers *CDH1*, *CD24*, and *EpCAM*. *ALDH1A1*^+^ cells also expressed significantly higher levels of progesterone receptor (*PGR*), androgen receptor (*AR*) and the ATP-binding cassette transporters *ABCG2* and *ABCB1* (**Supplemental Table 6**). These results suggest that *ALDH1A1* expressing cells are a distinct subpopulation with a mesenchymal phenotype that overexpress *PGR* and *AR* mRNA relative to the remainder of ALDH^+^ cells

### Clinical Relevance of ALDH^+^ Associated Genes in Breast Cancer

In light of our findings that within the population of ALDH+ cells, those cells that expressed both mesenchymal and epithelial genes had the greatest mammosphere forming capacity, we determined whether the gene signature of cells with these properties (cluster 2) had clinical relevance for women with breast cancer. We identified 4 genes that were significantly overexpressed in Cluster 2 cells: *KRT7*, *NOTCH3, CD146,* and *YAP1*, and analyzed publicly available gene expression data to access the clinical relevance of these genes in breast cancer (Figure 6A-D). These genes were overexpressed in triple negative breast cancers compared to other breast cancers in the Cancer Genome Atlas. This is consistent with previous reports for enrichment of CSC in this subtype (Idowu et al., 2012; Yoshioka et al., 2011). Furthermore, overexpression of each of these four genes was also associated with worse survival in triple negative breast cancer patients, and patients with high mean expression of all 4 genes had significantly worse survival (Figure 6E, HR = 2.22). These results suggest that gene expression programs involved in the regulation of Cluster 2 ALDH^+^ cells are dysregulated in the most aggressive triple negative breast cancers.

**Figure 6.**
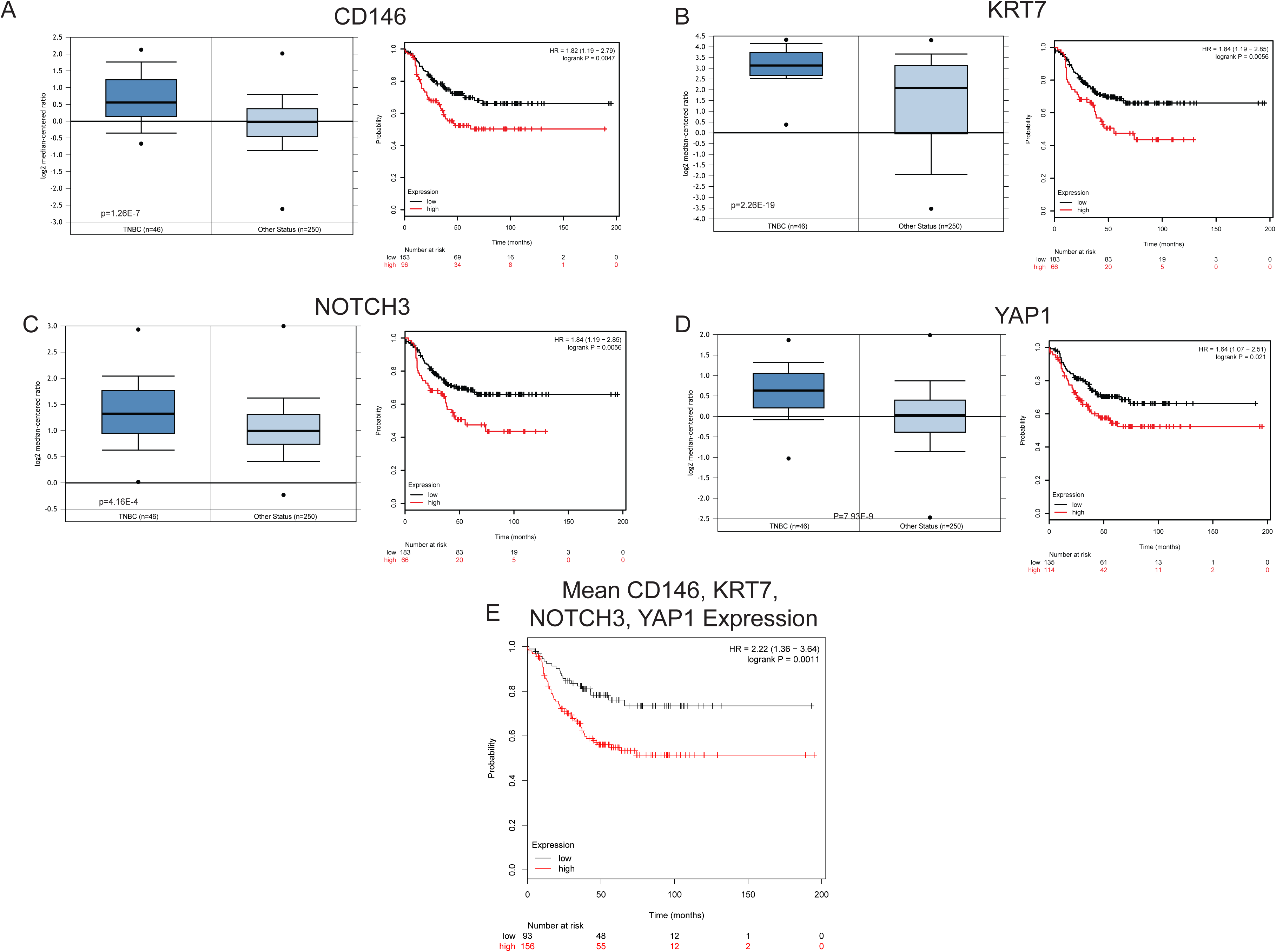
Expression of Cluster 4 genes in triple negative breast cancers (TNBCs) and the relationship of expression with survival. (A-D) Expression of *CD146, KRT7, NOTCH3, and YAP1,* respectively, in TNBCs relative to other breast cancers as well as relative survival of patients with TNBC tumors that express high relative to low amounts of each gene. (E) Relative survival of TNBC patients with tumors that have high relative to low mean *CD146, KRT7, NOTCH3, and YAP1* expression.

## Discussion

The results presented here highlight the inter- and intraindividual heterogeneity of the stem/progenitor cell populations of the normal human mammary gland. By conducting high throughput RNA sequencing on flow cytometry sorted ALDH^+^ and CD44^+^ cells, we identified significant differences, as well as similarities, between these two populations when compared to differentiated CD24^+^ cells. CD44^+^ express high levels of mesenchymal-related markers, while ALDH^+^ cells, in bulk, express high levels of epithelial-associated markers, but also express intermediate levels of some mesenchymal-associated markers. We identified a considerable amount of variation in the proportion of ALDH^+^ cells between individuals, and further, substantial variation between individuals when ALDH^+^ cells were subsequently analyzed for their expression of CD44 and CD24. Cells that express both ALDH and CD44^+^CD24^−^ have the highest mammosphere formation potential. At the single cell level, compared to the rest of ALDH^+^ cells, ALDH^+^CD44^+^CD24^−^ cells overexpress EMT and stemness genes, including *ID1, TWIST1*, and *SOX2*. By unbiased clustering of ALDH^+^ single cell RNA expression data, we identified substantial heterogeneity within this population, including individual cells with epithelial-like, mesenchymal-like, or dual epithelial/mesenchymal phenotypes. Within the cells that expressed both epithelial and mesenchymal markers, there was high expression of genes which are overexpressed in triple negative breast cancer and are associated with poor survival in these patients.

Our previous work has shown that breast cancer stem cells express the markers CD44^+^CD24^−^ (Al-Hajj et al., 2003) or ALDH^+^ (Ginestier et al., 2007), and that these two marker sets isolate distinct populations of mesenchymal-like and epithelial-like cells, respectively (Liu et al., 2013). Despite the lack of overlap between these populations, these cell maintain a cellular plasticity allowing them to inter-convert in a process regulated by the tumor microenvironment (Liu et al., 2013). Similarly, we previously showed that ALDH^+^ or CD44^+^ normal mammary cells cultured as mammospheres also have distinct epithelial or mesenchymal phenotypes relative to each other (Colacino et al., 2016). In the current study, we expanded upon these observations by using CD24^+^ cells as a differentiated cell anchor to compare the gene expression signatures of ALDH^+^ and CD44^+^ cells. We determined that CD44^+^ cells are strongly enriched for the mammary stem cell gene signature of CD49f^+^/EpCAM- cells (Lim et al., 2009). In contrast, there was no clear enrichment for either the mammary stem cell or the luminal progenitor gene signature in ALDH^+^ cells. Despite large gene expression and phenotypic differences between CD44^+^ and ALDH^+^ cells, when compared to CD24^+^ cells, there was substantial overlap in differentially expressed biological pathways, including the proteasome, oxidative phosphorylation, focal adhesion, and ECM-receptor interactions. ALDH^+^ and CD44+ cells displayed similar patterns of expression of genes associated with the proteasome and oxidative phosphorylation. Findings of increased proteasome activity in these two stem/progenitor cell enriched populations corroborate previous findings in embryonic stem cells (Jang et al., 2014; Vilchez et al., 2012) and neural progenitor cells (Zhao et al., 2016). Intriguingly, there are reports that breast cancer stem cells express lower proteosomal activity (Vlashi et al., 2009; Vlashi et al., 2013), suggesting that downregulation of the proteasome may be a key step in the generation of breast cancer stem cells. Both ALDH^+^ and CD44^+^ cells also expressed higher levels of genes involved in oxidative phosphorylation. While it has been assumed that the stem cell niche is hypoxic and that stem cells preferentially utilize glycolysis for energy production (Shyh-Chang et al., 2013), emerging data also points to a number of proliferative undifferentiated cell populations utilizing oxidative phosphorylation, including osteoblasts (Chen et al., 2008) and pre-adipocytes (D’Esposito et al., 2016; Tormos et al., 2011). Finally, compared to CD24^+^ cells, both CD44^+^ and ALDH^+^ cells significantly differentially express genes associated with focal adhesion and interactions with the ECM, although the gene expression patterns between the two stem/progenitor enriched fractions are very different. ECM-interacting receptors and proteins, including cadherins and integrins, are known key regulations of stem cell self-renewal and differentiation (Gattazzo et al., 2014).These interactions may play an important role in formation of the stem cell niche. For example, *ITGB3* was expressed at higher levels in ALDH^+^CD44^+^CD24^−^ cells at the single cell level, and in CD44^+^, compared to CD24^+^, cells in sorted population analyses, suggesting that this integrin molecule may be associated with CD44 expression. Further exploration of the influence of these pathways in the regulation of normal breast stem cells will likely provide new key insights into breast cancer stem cell biology.

We identified substantial heterogeneity in proportions of ALDH^+^ stem cells between individuals. Moreover, there was considerable interindividual variation in CD44^+^CD24^−^ expression within the ALDH^+^ population. It is currently unknown what epidemiologic, clinical, or environmental risk factors contribute to this variation, although this is an area of research interest for us moving forward. The finding of a substantial normal ALDH^+^CD44^+^CD24^−^ (Figure 4A) population stands in contrast to our previous findings in cancer (Liu et al., 2013). Normal ALDH^+^CD44^+^CD24^−^ cells had the highest mammosphere forming potential, providing a functional readout of the stem/progenitor cell-like behavior of these cells. Interestingly, an analysis of the luminal portion of the human mammary gland identified that EpCAM^+^CD49f^+^ALDH^+^ cells were previously found to have no overlap with CD44^+^CD24^−^ (Shehata et al., 2012), while we identified clear overlaps between the ALDH^+^ and CD44^+^CD24^−^ population as well as a range of phenotypes from mesenchymal to epithelial within ALDH^+^ cells, illuminating the existence of ALDH^+^ populations outside of the luminal progenitor context (Eirew et al., 2012). ALDH1A3 was previously identified as the isoform with highest aldehyde dehydrogenase activity, by the Aldefluor assay, in the breast (Marcato et al., 2011), which we corroborated with our RNA-seq data of the bulk ALDH^+^ population. However, at the single cell level, we identified a small number of ALDH+ cells that expressed *ALDH1A1* and had mesenchymal phenotypes. Honeth and colleagues reported that ALDH1A1 and ALDH1A3 staining cells are exclusive from each other, with ALDH1A1 cells localizing in small lobules and ALDH1A3 cells localizing in the extralobular ducts (Honeth et al., 2015). The different localization and phenotype of ALDH1A1 and ALDH1A3 cells raises the possibility that cells expressing these markers have different functions or may be active during different timepoints in development. ALDH1A1 expression has also previously been shown to be important for mammosphere formation potential (Honeth et al., 2014), and normal breast tissues from women with BRCA1 mutations were enriched for ALDH1A1 positive cells (Honeth et al., 2015; Liu et al., 2008). It is also possible that, despite us depleting CD140b positive stromal cells, a few ALDH^+^ stromal cells were isolated for single cell analyses, as stromal breast cells express ALDH1A1 (Eirew et al., 2012). Future efforts should focus on the development of methods to distinguish and isolate live human ALDH1A3 and ALDH1A1 cells for further characterization and experimental interrogation.

The epithelial-to-mesenchymal transition (EMT), and the converse process, MET are essential in the formation of cancer metastases. Emerging evidence is now pointing to EMT and MET not as static states, but a continuum, with data showing that cells exhibit both phenotypes in both development and cancer (Nieto, 2013). Induction of EMT in non-tumorigenic human mammary epithelial cells by ectopic expression of the EMT transcription factors Twist or Snail is sufficient to generate CD44^+^CD24^−^ stem cells (Mani et al., 2008).Transient activation of the EMT transcription factor TWIST1 in mammary epithelial cells can induce a stem-like phenotype, without full reversion to an epithelial state (Schmidt et al., 2015). Furthermore, certain populations within circulating tumor cells display a dual epithelial/mesenchymal phenotype, which further underscores the importance of understanding the biology of this hybrid dynamic (Aceto et al., 2014). Single cell profiling of normal and cancerous breast cells showed that early circulating breast cancer cells have a phenotype that clearly resembles that of a normal breast stem cell (Lawson et al., 2015).

We identified a subpopulation of ALDH^+^ breast cells that were highly enriched for “stemness” genes that also expressed a hybrid of epithelial and mesenchymal markers. Additionally, genes overexpressed in this subpopulation were found to be overexpressed in TNBC and predictive of survival in TNBC as well, suggesting that the pathways regulating this subpopulation of ALDH cells are important in the most aggressive forms of breast cancer. Others have also begun to report on the properties of epithelial/mesenchymal hybrid cells in the breast. Culture of murine mammary epithelial EpH4 cells in an alginate 3D matrix and transiently treated with TGFβ1 lead to the development of epithelial/mesenchymal hybrid cells, which exhibited increased clonogenicity and a more invasive phenotype (Bidarra et al., 2016). Additionally, single cell qPCR analysis of CD44+ cells isolated from mammospheres derived from oncogene-immortalized human mammary epithelial cells identified a gene signature of a dual epithelial/mesenchymal phenotype (Grosse-Wilde et al., 2015). Mathematical modeling, validated with MCF10A *in vitro* cultures, predicts that Notch-Jagged signaling regulates the emergence of hybrid epithelial/mesenchymal cells (Boareto et al., 2016). We identified that our normal breast epithelial/mesenchymal cells were also enriched for members of the Notch signaling pathway. Our results show that epithelial/mesenchymal hybrid cells exist in the normal human breast. Further elucidation of the biological processes that regulate these cells will likely provide new targets for the prevention and treatment of metastatic cancer.

This study highlights the advantages of assaying sorted cell populations, and single cells, rather than bulk tissue from the normal human breast to provide new insights into mammary gland biology. From our RNA-seq analysis of ALDH^+^ cells, we observed that ALDH^+^ cells express genes associated with both epithelial and mesenchymal cell states. However, it was not until we quantified at single cell gene expression from FACS isolated ALDH^+^ cells that we observed that there are subsets of epithelial-like, mesenchymal-like, and hybrid epithelial/mesenchymal cells. Understanding the factors that regulate the relative proportions of these cells is likely important for our understanding of the origins of different breast cancer subtypes. Future work, taking advantage of new advances in single cell transcriptomics, such as Drop-Seq (Macosko et al., 2015), will undoubtedly reveal additional levels of complexity in human mammary stem cell regulation. Even with a more limited spectrum of genes analyzed a single breast stem cell transcriptional profile, our work reveals striking inter- and intraindividual heterogeneity in human mammary stem cell populations and identifies genes and pathways that help define this heterogeneity. Further understanding of the intrinsic and extrinsic factors that impact breast stem cell heterogeneity will likely provide key insights for the prevention, early detection, and treatment of breast cancer.

## Acknowledgements

Support for this study was provided by grants from the National Cancer Institute (R03 CA167700), the National Institute of Environmental Health Sciences (T32 ES007062) and the National Human Genome Research Institute (NHGRI) (T32 HG00040).

